# Combining the 5.8S and ITS2 gene regions to improve classification of fungi

**DOI:** 10.1101/532358

**Authors:** Felix Heeger, Christian Wurzbacher, Elizabeth C. Bourne, Camila J. Mazzoni, Michael T. Monaghan

**Author notes:** these authors contributed equally to this work (Camila J. Mazzoni, Michael T. Monaghan).

## Abstract

- The internal transcribed spacer (ITS) is used in DNA metabarcoding of fungi. One disadvantage of its high variability may be a failure to classify OTUs when no similar reference sequence exists. We tested whether the 5.8S region, often sequenced with ITS2 but discarded before analysis, could provide OUT classifications when ITS fails.
- We used *in silico* evaluation to compare classification success of 5.8S and ITS from the UNITE database when reference sequences of the same species, genus, or family were removed. We then developed an automated pipeline for a combined 5.8S - ITS2 analysis and applied it to mixed environmental samples containing many lineages that are underrepresented in databases.
- ITS was clearly superior for species-level classifications with a complete reference database, but 5.8S outperformed ITS at higher level classifications with an incomplete database. Our combined 5.8S-ITS2 pipeline classified 3x more fungal OTUs compared to ITS2 alone, particularly within Chytridiomycota (10x) and Rozellamycota (3x).
- Missing reference sequences led to the failure of ITS to classify many fungal OTUs at all, and to a significant underestimation of environmental fungal diversity. Using 5.8S to complement ITS classification will likely provide better estimates of diversity in lineages for which database coverage is poor.

## Introduction

The Fungi comprise an enormous diversity of species and life styles. Estimations of the number of species range from 2.2 to 3.8 million (Hawksworth & Lücking, 2017) of which only a small fraction (<145,000, http://www.speciesfungorum.org/Names/Names.asp, accessed January 2019) have been formally described. The evolutionary relationships between major fungal lineages are far from resolved an there is still no general agreement on the number of phyla, particularly for the basal clades. Hibbett et al. (2007) named seven phyla. Blackwell (2011) gave the number of phyla as “about 10”. Following the recent definition of Rozellomycota (or Cryptomycota) (Lara *et al.*, 2010; Jones *et al.*, 2011; Corsaro *et al.*, 2014), Tedersoo *et al.* (2017b) mentions 12 phyla and indicates that there may be more phyla. The latest taxonomy defines 16 basal phyla in addition to the Ascomycota and Basidiomycota bringing the total to 18 (Wijayawardene *et al.*, 2018). The community-curated reference database UNITE (Kõljalg *et al.*, 2013) currently (version 7.2, 2017-12-01) also lists 18 phyla, including the preliminary named phyla GS01 and GS19.

Schoch *et al.* (Schoch *et al.*, 2012) proposed the internal transcribed spacer (ITS) region of the eukaryotic rRNA operon as a universal fungal DNA barcode. The ITS region is ca. 300- 1,200 bp and is located between the 18S (SSU) and 28S (LSU) rRNA genes. It contains the two highly variable spacers, ITS1 and ITS2, separated by the less variable 5.8S gene (Nilsson *et al.*, 2008). The full ITS region is included in the UNITE database (Kõljalg *et al.*, 2013).

Advances in sequencing technologies have enabled a shift to DNA metabarcoding surveys of environmental samples, whereby sample throughput is much higher than previously possible and whole communities can be studied without the need for isolation and culture of single species or isolation of genotypes through cloning of single DNA fragments (Nilsson *et al.*, 2018). Because the maximum length of continuously read sequence (~550 bp with overlapping paired-end design) using the most commonly used Illumina sequencer for metabarcoding (MiSeq), it is not feasible to sequence the whole ITS region. Most studies focus on either the ITS1 or ITS2 (Tedersoo *et al.*, 2014; Miller *et al.*, 2016; Wurzbacher *et al.*, 2017) as a result.

The ability of Illumina based DNA metabarcoding to identify fungal taxa in mixed samples varies among studies. An *in silico* test with 8967 ITS sequences from a range of fungal phyla (Porras-Alfaro *et al.*, 2014) reported that > 90% of test data (ITS1 91%; ITS2 93%) were identified to the correct genus. In a mock community of 24 Dikarya species, both ITS1 and ITS2 sequences of different species could be clustered into one operational taxonomic unit (OTU) each and classified correctly (Tedersoo *et al.*, 2015). In environmental samples classification of ITS sequences has proven more challenging in many studies. Rime *et al.* (2015) reported that 5% of the ITS2 OTUs from soil samples could not be classified to phylum (i.e. only to kingdom fungi). Wurzbacher *et al.* (2017) found that 25% of fungal OTUs in permafrost thaw ponds could not be assigned to phylum with ITS2. In a study of fungi in decaying wood Yang *et al.* (2016) found that 19 - 25% of OTUs could not be classified below kingdom level, and a study from lake sediments reported 72% of fungi were unclassified for the ITS1 region and 49% of unclassified fungi for the ITS2 (Wahl *et al.*, 2018). These results all highlight the fact that the incomplete state of reference databases for many fungal taxa may hinder ITS classification, although it is not clear which taxonomic levels are affected and how this affects classification success.

A potential reason for the failure of ITS to classify fungal OTUs from environmental samples, even to higher taxonomic levels, is the variability of the ITS sequence itself. While high variability among closely related taxa makes the ITS an excellent DNA barcode, its variability also hinders classification of evolutionarily more distant taxa. This is because large sequence divergence can make it difficult to establish homology and impairs an alignment to identify a sister taxon. This may be especially problematic in less well studied habitats such as freshwater, where a wide variety of early diverging fungal lineages occur (Grossart *et al.*, 2016; Rojas-Jimenez *et al.*, 2017) and for which sequences from closely related species are often not available in reference databases.

Interestingly, many fungal DNA metabarcoding studies amplify the ITS2 region using the primer pair ITS3/ITS4 (White *et al.*, 1990), which includes a ~130 bp long fragment of the 5.8S rRNA gene that is normally discarded during the amplicon data processing steps (e.g. Lindahl *et al.*, 2013; Bálint *et al.*, 2014). The 5.8S rRNA gene has a much lower substitution rate compared to ITS1 or ITS2 sequence (Nilsson *et al.*, 2008) and is thus usually neglected as a potential barcode, but has been used for phylogenetic classification (Roose-Amsaleg *et al.*, 2004; Neubert *et al.*, 2006). The fact that the 5.8S gene is included in the full ITS reference database UNITE, allows for direct taxonomic comparison with the ITS1 and ITS2.

Here we tested whether the more conserved 5.8S region could provide higher level classification of fungi in cases where ITS2 could not, using *in silico* analysis of sequences present in the UNITE database. We classified query sequences at different taxonomic ranks using the 5.8S, ITS1 and ITS2 and examined the extent to which the classification success depended on database completeness. Specifically, we excluded all other sequences from individuals of either the same species, genus, or family. We observed that ITS1 and ITS2 are clearly superior for species-level classifications when the reference database is complete, but that 5.8S outperforms both markers at higher level taxonomic classifications with an incomplete database. Based on this result, we developed and implemented an automated pipeline to analyze amplicons that contain both 5.8S and ITS2 rRNA gene regions, typical of many fungal DNA metabarcoding studies. A test on sequence data from sediment and water samples from 20 freshwater lakes showed that the 5.8S sequence added phylum level classifications for most (74%) of the 64% of our ITS2 OTUs that were unclassified with ITS2 alone. The current version of the pipeline can be found at www.github.com/f-heeger/two_marker_metabarcoding.

## Material and Methods

### Testing the effects of an incomplete reference database

For the *in silico* evaluation of how database completeness affects classification with different rRNA markers, we created a dataset whereby the classification of each query sequence was known, and where at least one other sequence from (a) the same species, (b) a different species in the same genus, and (c) a different genus within the same family, were also available. This allowed us to test whether classifications at a given rank were correct, even when all other sequences for the species, genus, or family were not present (i.e. removed from our reference database). An additional criterion was that complete sequences of ITS1, ITS2, and 5.8S had to be available to allow for comparison between the markers. We created such a dataset in the following way: Fungal ITS1, 5.8S and ITS2 sequences were extracted from sequences in the UNITE database (version 7.2, 2017-12-01) using ITSx with default parameters (version 1.0.11, Bengtsson-Palme *et al.*, 2013). Sequences that satisfied the following three criteria were selected: i) all three markers could be detected by ITSx, ii) a species-level classification was available in UNITE, and iii) at least one other sequence was available from the same species, from the same genus (but different species), and from the same family (but different genus). There were 5038 sequences that satisfied these criteria and from these we chose a random subset of 100 sequences for our evaluation.

Marker sequences (ITS1, ITS2, 5.8S) were classified independently with the lowest common ancestor (LCA) classification approach based on database search results similar to the one employed in MEGAN (Huson *et al.*, 2007). First a database search of each sequence is performed against the UNITE database. For each sequence hits with an e-value below a minimum value (default: 10^−7^) are considered. Any hit with an identity or query coverage below a certain threshold (default: 80% and 85% respectively) or a bitscore lower than a certain percentage (default: 95%) of the best score for that sequence is excluded. For the remaining hits the lowest common ancestor in the taxonomic tree that underlies UNITE is determined in the following way: For each level in the taxonomic tree, starting from kingdom, classifications of all hits are compared. If the classification of a certain percentage (default: 90%) or more of the hits at this taxonomic level are the same, it will be accepted as the classification on this level for the query sequence. Otherwise the lowest common ancestor is found and the query will only be classified to the last level, where a majority was achieved. During this process any classifications of “undetermined” or “unclassified” are ignored.

ITS2 sequences were additionally analyzed with the RDP (Wang *et al.*, 2007) classifier to make sure that the LCA approach we implemented here gives results comparable to widely applied tools. We employed the classifier trained for use in the PIPITS pipeline (Gweon *et al.*, 2015) on ITS sequences from the current version (7.2, 2017-12-01) of UNITE.

For 5.8S and ITS2, the classification was run using a range of parameter values for minimum identity, minimum coverage, top bit score fraction cutoff, and LCA majority stringency. This was done to investigate the parameter stability of the approach. The effect of missing database coverage was tested by first classifying query sequences using the complete reference database, and then repeating the process three times, removing all sequences from the same species, genus, and family in subsequent iterations. To assess whether classifying the 5.8S and ITS2 together was an effective method, we classified the combined 5.8S and ITS2 fragment with the LCA approach and compared the resulting classifications with those in the UNITE database.

### 5.8S reference data set

As a reference dataset for classification of 5.8S sequences, we used the 5.8S sequences that were extracted from UNITE with ITSx (above) and complemented them with non-fungal 5.8S sequences from the 5.8S rRNA family (RF00002) of the Rfam database (Kalvari *et al.*, 2018). Identical sequences were dereplicated to one representative with vsearch (Rognes *et al.*, 2016). For each representative sequence, a taxonomic classification was determined by generating a LCA from the classifications of all sequences it represents. For Rfam sequences classified as fungi, any classification at lower rank was ignored and priority was given to the taxonomy information from the UNITE database.

### Description of the pipeline

The pipeline was implemented as a workflow with snakemake (Köster & Rahmann, 2012) with four main stages: 1) initial read processing, 2) 5.8S classification, 3) ITS2 classification and 4) final classification (Fig. 1).

1. Initial read processing starts by producing quality plots with FastQC (version 0.11.2, Andrews). The presence of the forward or reverse primer in the first 25 bp of the respective read is checked with flexbar (version v2.5_beta, Roehr *et al.*, 2017). Quality trimming with Trimmomatic (version 0.35, Bolger *et al.*, 2014) consists of a sliding window trimming (default window size: 8 and a minimum Phred score: 20) and removal of trailing bases with a low (default: <20) Phred quality, followed by the removal of sequences that are too short (default: < 200) or have a low average Phred quality (default: <30) after trimming. Forward and reverse reads of each pair are then merged with Pear (version 0.9.6, Zhang *et al.*, 2014). By default the minimum overlap for merging is set to 10. Pairs that cannot be merged or are too short (default: < 150) or too long (default: > 550) after merging are discarded. Merged sequences are dereplicated with vsearch. Potential chimeras (including sequences classified as “suspicious”) are removed with vsearch in *de novo* chimera detection mode with default parameters. The 5.8S and ITS2 sequences are extracted with ITSx with default parameters, except that partial 5.8S sequences are accepted. The 5.8S and the ITS2 sequences are independently classified in stage 2 and 3 respectively.
2. 5.8S classification starts with removal of the forward primer and sequences with ambiguous bases are discarded using cutadapt (version 1.9.1, Martin, 2011). Sequences are dereplicated with vsearch and then classified by a similarity search against our combined 5.8S reference dataset (above) with lambda (version 0.9.3, Hauswedell *et al.*, 2014) followed by a LCA classification as described for the *in silico* test (above).
3. ITS2 classification starts with dereplication of ITS2 sequences with vsearch. Clustering into OTUs is done with swarm2 (version 2.1.6, Mahé *et al.*, 2015). OTUs are classified by similarity search and LCA in the same way as 5.8S sequences are classified (above).
4. The final classification combines the classifications from stage 2 and 3. For each read present in an ITS2 OTU cluster, all 5.8S sequences and their classifications are collected. The 5.8S classifications are combined with the same LCA approach explained above. The resulting classification is compared to the ITS2 classification. If 5.8S and ITS2 classification are concordant, but the ITS2 is classified to a lower taxonomic rank, the ITS2 classification is accepted. Sequences that are unclassified with ITS2 will automatically take the 5.8S classifications. All conflicting classifications can either be marked (default) or resolved by the user by giving priority to one of the markers.

**Figure 1:**
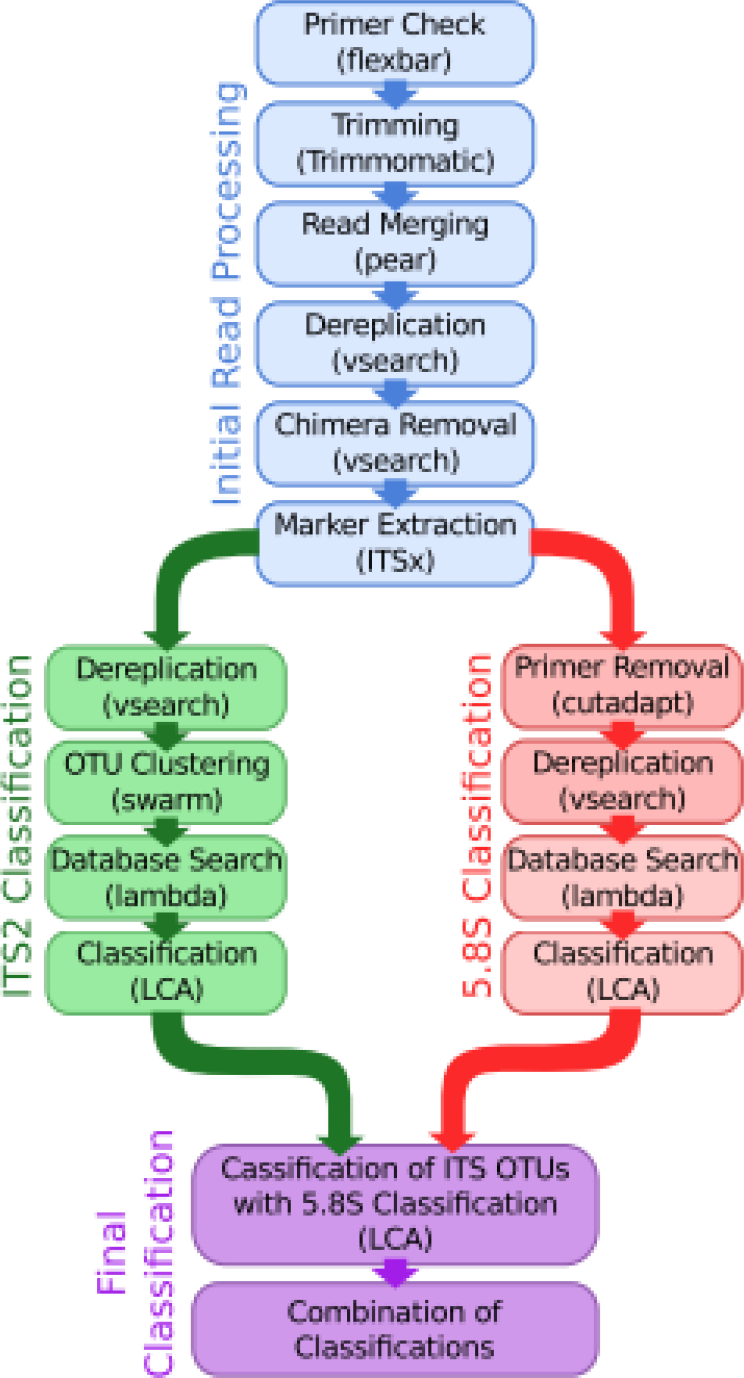
Overview of the steps in the automated pipeline for parallel classification with ITS2 and 5.8S. External tools and approaches used are given in parentheses.

### Test with reads from freshwater lake samples

We tested the pipeline on an unpublished data set (Bourne, E.C. *et al.* unpublished) of water and sediment samples, taken in October and November 2014 from the littoral zone of 20 freshwater lakes in North-East Germany. In six lakes, additional sediment and water samples were taken from the pelagic zone. Amplification was performed using ITS3mix1 and ITS3mix2 forward primers (Tedersoo *et al.*, 2015) that were modified by adding a degenerate base (W) at the third position, and the standard reverse primer ITS4 (*White et al.*, 1990). This primer set amplified a 350-500 bp amplicon consisting of the full ITS2 and ca. 130 bp of the 5′-end of the 5.8S gene. Amplicons were sequenced with overlapping 300 bp paired-end reads on an Illumina MiSeq (v3 chemistry).

## Results

Analysis of the classification of query sequences with an increasingly incomplete reference database showed a clear difference among markers (Fig. 2). When the query species was present in the reference database, ITS1 and ITS2 both correctly classified 90% of queries to species, whereas 5.8S classified 3% of queries to species and 56% of sequences to order (Fig 2). The removal of all sequences from the same species, genus, or family had an increasingly detrimental effect on the classification success of both ITS sequences (Fig. 2). Removing only the query species (i.e. other species in the genus still present in the database) caused a distinct drop in successful classification of ITS1 and ITS2 at the kingdom (from 100% to 83% and 88% respectively), phylum (from 100% to 83% and 88% respectively), and class (from 100% to 83% and 87% respectively) ranks (Fig. 2). In contrast, the kingdom and phylum rank classifications of 5.8S sequences were not notably affected by the removal of reference sequences, with classification at the class rank only dropping from 83% to 81% and classification to kingdom and phylum being completely unaffected (Fig. 2).

**Figure 2:**
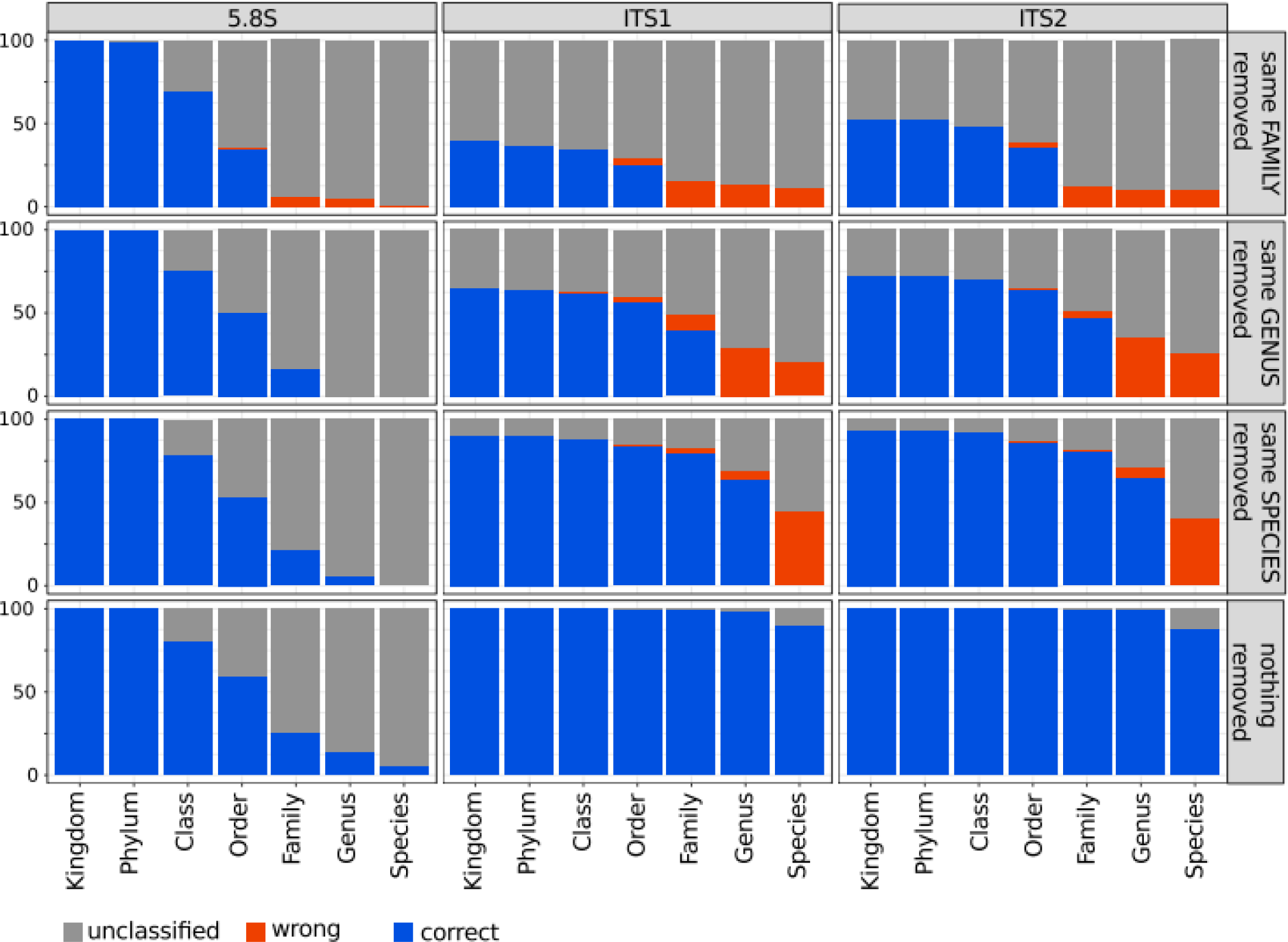
Results from the classification of the *in silico* test set (100 sequences). LCA classification was performed with different markers (panels from left to right) and with different completeness of the reference database (panels from top to bottom). Numbers of correct (blue), wrong (red) and unassigned (grey) classifications are given compared to the original classification in UNITE.

The LCA classification was performed with different parameters for ITS2 and 5.8S to test parameter stability. The stringency parameter had minimal influence on ITS2 classifications (Fig. S1). Lowering the parameters of minimum identity (Fig. S2) and minimum coverage (Fig. S3) increased the number of classifications, but also increased the numbers of wrong classifications. Lower values for the top bitscore fraction parameter caused more wrong ITS2 classifications without increasing the number of correct classifications (Fig. S4). Minimum identity and minimum coverage had little influence on 5.8S classifications (Fig. S5 and S6), although a very high value (100%) resulted in more wrong classifications. The top bitscore fraction parameter gave more correct 5.8S classifications for values ≤ 5%, but at the cost of an increased number of wrong classifications (Fig. S7). Finally a low value (≤ 85%) for the stringency parameter resulted in more wrong 5.8S classifications, while a very high value (100%) led to a decrease in correct assignments (Fig. S8).

Comparison with RDP classifications (Fig. S9) showed that the LCA approach gives comparable results to the RDP classifier (trained on the UNITE database) for our data. The comparison between independent classification of ITS2 and 5.8S with the classification of a combined fragment of both regions revealed that a combined fragment improved classification at kingdom and phylum ranks, but not to the same extent as an independent classification of 5.8S and ITS2 with a subsequent combination of the result (Fig. S10).

The environmental data set from 20 freshwater lakes (water and sediment samples) consisted of 13.6 million read pairs. Our analysis pipeline generated 17,514 non-singleton OTUs. The 5.8S marker classified nearly three times as many OTUs compared to ITS2, including a 28-fold increase in the number of Chytridiomycota OTUs and a 6-fold increase in Rozellomycota TOUs (Tab. 1). Using ITS2, 30% of all OTUs were classified as Fungi, 1% were classified as belonging to a different kingdom, and 69% were unclassified (Fig 3). In contrast, using 5.8S, 64% were classified as Fungi, 12% were classified as belonging to a different kingdom, and 24% were unclassified (Fig. 3). Using the two markers in combination, results were very similar to those using 5.8S alone (Tab. 1), but with more low level (family to species) classifications (Fig. 3).

**Table 1.** Water and sediment OTU classification (17514 OTUs, based on ITS2 clustering) using ITS2 and 5.8S markers individually and in combination.

|  | Unclassified | Fungi | Ascomycota | Basidiomycota | Chytridiomycota | Rozellomycota | Other Fungal<br>Phyla | Non-Fungi |
| --- | --- | --- | --- | --- | --- | --- | --- | --- |
| ITS2 | 12071 | 5262 | 2983 | 1946 | 111 | 60 | 68 | 181 |
| 5.8S | 4123 | 11263 | 3649 | 2439 | 3030 | 339 | 215 | 2128 |
| ITS2 + 5.8S | 4107 | 11262 | 3651 | 2441 | 3031 | 339 | 215 | 2145 |

**Figure 3:**
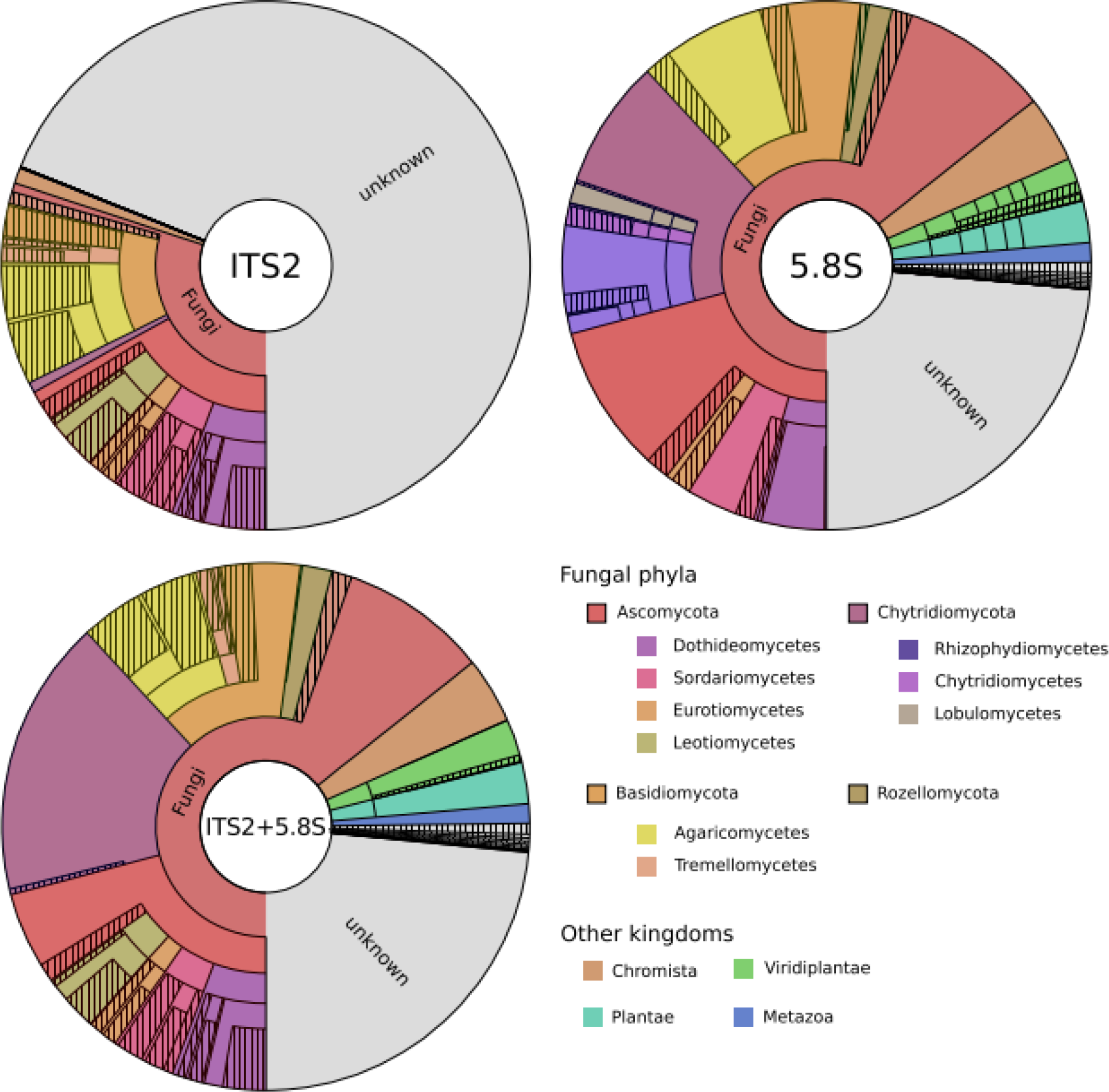
Classification of OTUs from lake water and sediments when using ITS2 (top left) or 5.8S (top right) and combined classification with our pipeline (bottom left). Concentric circles from the inside out represent levels of taxonomic classification from kingdom to species. Hatched areas contain more specific classifications that are not shown. Segments are colored by kingdoms and for fungi by phyla and classes.

There was a classification conflict for only one OTU. The 5.8S classification was Arthropoda, whereas the ITS2 classification was Ascomycota. This was caused by a miss-classification of SH200261.07FU in the UNITE database (R.H. Nilsson pers. comm.), that has been subsequently corrected in UNITE.

## Discussion

We developed and implemented a modular pipeline for the processing of fungal DNA metabarcoding data that uses the taxonomic information from the 5.8S gene to complement the more widely used ITS2 region. These markers are adjacent to one another in the eukaryotic rRNA operon and >100 bp of 5.8S are typically sequenced using the most frequently employed ITS2 primer sets (White *et al.*, 1990; Tedersoo *et al.*, 2015), but then discarded prior to analysis. Using both markers in combination allowed us to classify a substantially greater number of OTUs than with ITS2 alone, in particular for less well studied, basal fungal lineages.

Our *in silico* analysis of the UNITE database expanded on earlier results (Porras-Alfaro *et al.*, 2014) that ITS1 and ITS2 are very good marker sequences when the database contains the exact same sequence or at least a sequence from the same species. In our test cases, no sequences were assigned to the wrong species and very few were unclassified. However, when only removing all sequences of the same species from our reference dataset, the ability to classify the genus dropped to 71% and 70% for ITS1 and ITS2 respectively, despite there being representatives of the genus in the reference dataset. Even for higher taxonomic ranks (phylum, class) the removal of the species caused classification problems. Simulating novel genera or families by removing the respective sequences from the database increased the effect even more. This is most likely the reason that many fungal OTUs remain unclassified in environmental studies that focus on poorly studied environments like freshwater (Grossart *et al.*, 2016; Rojas-Jimenez *et al.*, 2017). We found that new species, genera or families that do not have any reference sequences available are often unidentified at even at the kingdom rank, leading to fungal diversity being severely underestimated.

In our environmental data set from lake water and sediments, there were large differences in OTU classifications, depending on whether we used ITS2 or 5.8S. The proportion of OTUs that could be identified as fungi was twice as high using the 5.8S., with 30-fold more Rozellomycota (also known as Cryptomycota) and nearly 10-fold more Chytridiomycota. Chytridiomycota are not well represented in the UNITE database (Frenken *et al.*, 2017) and our *in silico* analysis showed that even if our environmental OTUs were represented by other members of the same genus or family, the ITS2 classification can fail completely. As a result, using ITS2 alone would have led to an estimate of Chytridoimycota of 3%, while the 5.8S classifications indicate that the actual proportion is an order of magnitude higher (32%). Similarly, the percentage of Rozellomycota would increase from 0.1% to 3% (Fig. 3). An estimation of the proportion of fungal phyla based on the ITS2 alone would have been strongly biased towards Ascomycota and Basidiomycota, which are better represented in the reference database.

Although the ITS2 barcode allows for accurate identification when near-perfect reference data are available, it may be unable to find a high enough similarity to any sequence when no closely related species is represented in the database. In such cases, the 5.8S sequence can help to classify OTUs to at least a higher taxonomic rank. In our environmental data, the 5.8S was especially helpful in splitting the results into fungal and non-fungal sequences when it comes to early diverging lineages or lineages that belong to the Top 50 unknown fungal lineages (Nilsson *et al.*, 2016). Nonetheless, our results clearly indicate that the 5.8S would be of limited use as a DNA barcode on its own, or to delineate OTUs, but it should rather be seen as providing complementary information.

Our implementation of LCA-based classification performed comparably to the commonly used RDP classifier on our test dataset and was not very sensitive to parameter choice. This indicates that our implementation is working as well as commonly used approaches and can be used to study the advantage of using multiple markers as well as the influence of an incomplete database. Unlike using a single “best” (e.g. lowest e-value) blast hit for identification which is problematic due to stochastic ranking of top hits (Shah *et al.*, 2018) and can easily lead to wrong classifications if the query species is missing from the database, our approach uses a certain proportion of top blast hits to try and quantify the uncertainty of our classifications by choosing a higher taxonomic rank. Nevertheless we found a substantial amount of wrong assignments in the *in silico* analysis, when the database was not complete (Fig. 2).

Third generation sequencing technologies currently available allow for the sequencing of longer amplicons. These include studies of the full-length 16S for bacteria (Mosher *et al.*, 2014; Schloss *et al.*, 2016; Singer *et al.*, 2016), the full ITS region (Schlaeppi *et al.*, 2016; Tedersoo *et al.*, 2017a) and most of the rRNA operon (Heeger *et al.*, 2018). Longer amplicons with multiple gene regions could be analyzed using the approach we have developed here. Although longer amplicons can increase identification success (Tedersoo *et al.*, 2017a), they typically result in lower sequencing depth because of the higher cost per base and can therefore increase the risk of missing rare taxa (Kennedy *et al.*, 2018). Primer pairs to target longer amplicons have also not yet been optimized to prevent primer and long-range amplification bias (Heeger *et al.*, 2018). We suggest that explicitly including the partial 5.8S gene into the analysis of shorter amplicons (as used in second-generation sequencing technologies such as applied here) can dramatically improve the high level classification of new species and poorly studied clades without increasing cost or reducing read depth.

## Acknowledgements

We thank R. Henrik Nilsson and Kessy Abarenkov for help with an earlier version of the manuscript, Susan Mbedi and Kirsten Richter for help with sequencing, and Hannah Ebbinghaus for help with PCRs. Research was partially funded by the Leibniz Association Pakt/SAW project “MycoLink” (SAW-2014-IGB-1).

## Author contribution

F.H., E.C.B., C.W., C.J.M. and M.T.M. conceived and designed the overall study. F.H. and
C.J.M. designed the analysis pipeline. F.H. implemented the analysis pipeline and carried out analysis. F.H., C.W., C.J.M. and M.T.M. wrote the manuscript and all authors contributed to the final manuscript. M.T.M. and C.J.M. contributed equally to this work.

## Supporting Information

**Fig. S1** Classification accuracy with ITS2 at different taxonomic levels with different values for the stringency parameter.

**Fig. S2** Classification accuracy with ITS2 at different taxonomic levels with different values for the minimum identity parameter.

**Fig. S3** Classification accuracy with ITS2 at different taxonomic levels with different values for the minimum coverage parameter.

**Fig. S4** Classification accuracy with ITS2 at different taxonomic levels with different values for the top-percent parameter.

**Fig. S5** Classification accuracy with 5.8S at different taxonomic levels with different values for the identity parameter.

**Fig. S6** Classification accuracy with 5.8S at different taxonomic levels with different values for the minimum coverage parameter.

**Fig. S7** Classification accuracy with 5.8S at different taxonomic levels with different values for the top-percent parameter.

**Fig. S8** Classification accuracy with 5.8S at different taxonomic levels with different values for the stringency parameter.

**Fig. S9** Classification accuracy with ITS2 with the LCA approach used in this article, the RDP classifier trained on the UNITE database (RDP_U), and the RDP classifier trained on the Warcup dataset (RDP_W).

**Fig. S10** Classification accuracy with the combined sequence of 5.8S and ITS2 (left), with 5.8S alone (middle), and with ITS2 alone (right) at different taxonomic levels and with different levels of database completeness.

## References

Andrews S. FastQC A Quality Control tool for High Throughput Sequence Data. http://www.bioinformatics.babraham.ac.uk/projects/fastqc/.

Bálint M, Schmidt P-A, Sharma R, Thines M, Schmitt I. 2014. An Illumina metabarcoding pipeline for fungi. Ecology and Evolution 4: 2642–2653.

Bengtsson-Palme J, Ryberg Martin, Hartmann Martin, Branco Sara, Wang Zheng, Godhe Anna, Wit Pierre, Sánchez García Marisol, Ebersberger Ingo, Sousa Filipe, et al. 2013. Improved software detection and extraction of ITS1 and ITS2 from ribosomal ITS sequences of fungi and other eukaryotes for analysis of environmental sequencing data. Methods in Ecology and Evolution 4: 914–919.

Blackwell M. 2011. The Fungi: 1, 2, 3 … 5.1 million species? American Journal of Botany 98: 426–438.

Bolger AM, Lohse M, Usadel B. 2014. Trimmomatic: a flexible trimmer for Illumina sequence data. Bioinformatics 30: 2114–2120.

Corsaro D, Walochnik J, Venditti D, Steinmann J, Müller K-D, Michel R. 2014. Microsporidia-like parasites of amoebae belong to the early fungal lineage Rozellomycota. Parasitology Research 113: 1909–1918.

Frenken T, Alacid Elisabet, Berger Stella A., Bourne Elizabeth C., Gerphagnon Mélanie, Grossart Hans Peter, Gsell Alena S., Ibelings Bas W., Kagami Maiko, Küpper Frithjof C., et al. 2017. Integrating chytrid fungal parasites into plankton ecology: research gaps and needs. Environmental Microbiology 19: 3802–3822.

Grossart H-P, Wurzbacher C, James TY, Kagami M. 2016. Discovery of dark matter fungi in aquatic ecosystems demands a reappraisal of the phylogeny and ecology of zoosporic fungi. Fungal Ecology 19: 28–38.

Gweon HS, Oliver Anna, Taylor Joanne, Booth Tim, Gibbs Melanie, Read Daniel S., Griffiths Robert I., Schonrogge Karsten, Bunce Michael. 2015. PIPITS: an automated pipeline for analyses of fungal internal transcribed spacer sequences from the Illumina sequencing platform. Methods in Ecology and Evolution 6: 973–980.

Hauswedell H, Singer J, Reinert K. 2014. Lambda: the local aligner for massive biological data. Bioinformatics (Oxford, England) 30: i349–355.

Hawksworth DL, Lücking R. 2017. Fungal Diversity Revisited: 2.2 to 3.8 Million Species. Microbiology Spectrum 5.

Heeger F, Bourne EC, Baschien C, Yurkov A, Bunk B, Spröer C, Overmann J, Mazzoni CJ, Monaghan MT. 2018. Long-read DNA metabarcoding of ribosomal RNA in the analysis of fungi from aquatic environments. Molecular Ecology Resources 0.

Hibbett DS, Binder M, Bischoff JF, Blackwell M, Cannon PF, Eriksson OE, Huhndorf S, James T, Kirk PM, Lücking R, et al. 2007. A higher-level phylogenetic classification of the Fungi. Mycological Research 111: 509–547.

Huson DH, Auch AF, Qi J, Schuster SC. 2007. MEGAN analysis of metagenomic data. Genome Research 17: 377–386.

Jones MDM, Richards TA, Hawksworth DL, Bass D. 2011. Validation and justification of the phylum name Cryptomycota phyl. nov. IMA fungus 2: 173–175.

Kalvari I, Argasinska J, Quinones-Olvera N, Nawrocki EP, Rivas E, Eddy SR, Bateman A, Finn RD, Petrov AI. 2018. Rfam 13.0: shifting to a genome-centric resource for non-coding RNA families. Nucleic Acids Research 46: D335–D342.

Kennedy PG, Cline LC, Song Z. 2018. Probing promise versus performance in longer read fungal metabarcoding. New Phytologist 217: 973–976.

Kõljalg U, Nilsson R. Henrik, Abarenkov Kessy, Tedersoo Leho, Taylor Andy F. S., Bahram Mohammad, Bates Scott T., Bruns Thomas D., Bengtsson Palme Johan, Callaghan Tony M., et al. 2013. Towards a unified paradigm for sequence based identification of fungi. Molecular Ecology 22: 5271–5277.

Köster J, Rahmann S. 2012. Snakemake—a scalable bioinformatics workflow engine. Bioinformatics 28: 2520–2522.

Lara E, Moreira D, López-García P. 2010. The Environmental Clade LKM11 and Rozella Form the Deepest Branching Clade of Fungi. Protist 161: 116–121.

Lindahl BD, Nilsson RH, Tedersoo L, Abarenkov K, Carlsen T, Kjøller R, Kõljalg U, Pennanen T, Rosendahl S, Stenlid J, et al. 2013. Fungal community analysis by high-throughput sequencing of amplified markers–a user’s guide. The New Phytologist 199: 288–299.

Mahé F, Rognes T, Quince C, Vargas C de, Dunthorn M. 2015. Swarm v2: highly-scalable and high-resolution amplicon clustering. PeerJ 3: e1420.

Martin M. 2011. Cutadapt removes adapter sequences from high-throughput sequencing reads. EMBnet.journal 17: 10–12.

Miller KE, Hopkins K, Inward DJG, Vogler AP. 2016. Metabarcoding of fungal communities associated with bark beetles. Ecology and Evolution 6: 1590–1600.

Mosher JJ, Bowman B, Bernberg EL, Shevchenko O, Kan J, Korlach J, Kaplan LA. 2014. Improved performance of the PacBio SMRT technology for 16S rDNA sequencing. Journal of Microbiological Methods 104: 59–60.

Neubert K, Mendgen K, Brinkmann H, Wirsel SGR. 2006. Only a Few Fungal Species Dominate Highly Diverse Mycofloras Associated with the Common Reed. Applied and Environmental Microbiology 72: 1118–1128.

Nilsson RH, Anslan S, Bahram M, Wurzbacher C, Baldrian P, Tedersoo L. 2018. Mycobiome diversity: high-throughput sequencing and identification of fungi. Nature Reviews Microbiology: 1.

Nilsson RH, Kristiansson E, Ryberg M, Hallenberg N, Larsson K-H. 2008. Intraspecific ITS Variability in the Kingdom Fungi as Expressed in the International Sequence Databases and Its Implications for Molecular Species Identification. Evolutionary Bioinformatics Online 4: 193–201.

Nilsson RH, Wurzbacher C, Bahram M, Coimbra VRM, Larsson E, Tedersoo L, Eriksson J, Duarte C, Svantesson S, Sánchez-García M, et al. 2016. Top 50 most wanted fungi. MycoKeys 12: 29–40.

Porras-Alfaro A, Liu K-L, Kuske CR, Xie G. 2014. From genus to phylum: large-subunit and internal transcribed spacer rRNA operon regions show similar classification accuracies influenced by database composition. Applied and Environmental Microbiology 80: 829–840.

Rime T, Hartmann Martin, Brunner Ivano, Widmer Franco, Zeyer Josef, Frey Beat. 2015. Vertical distribution of the soil microbiota along a successional gradient in a glacier forefield. Molecular Ecology 24: 1091–1108.

Roehr JT, Dieterich C, Reinert K. 2017. Flexbar 3.0 – SIMD and multicore parallelization. Bioinformatics 33: 2941–2942.

Rognes T, Flouri T, Nichols B, Quince C, Mahé F. 2016. VSEARCH: a versatile open source tool for metagenomics. PeerJ 4: e2584.

Rojas-Jimenez K, Wurzbacher C, Bourne EC, Chiuchiolo A, Priscu JC, Grossart H-P. 2017. Early diverging lineages within Cryptomycota and Chytridiomycota dominate the fungal communities in ice-covered lakes of the McMurdo Dry Valleys, Antarctica. Scientific Reports 7.

Roose-Amsaleg C, Brygoo Yves, Harry Myriam. 2004. Ascomycete diversity in soil feeding termite nests and soils from a tropical rainforest. Environmental Microbiology 6: 462–469.

Schlaeppi K, Bender SF, Mascher F, Russo G, Patrignani A, Camenzind T, Hempel S, Rillig MC, van der Heijden MGA. 2016. High-resolution community profiling of arbuscular mycorrhizal fungi. The New Phytologist 212: 780–791.

Schloss PD, Jenior ML, Koumpouras CC, Westcott SL, Highlander SK. 2016. Sequencing 16S rRNA gene fragments using the PacBio SMRT DNA sequencing system. PeerJ 4: e1869.

Schoch CL, Seifert KA, Huhndorf S, Robert V, Spouge JL, Levesque CA, Chen W, Consortium FB. 2012. Nuclear ribosomal internal transcribed spacer (ITS) region as a universal DNA barcode marker for Fungi. Proceedings of the National Academy of Sciences 109: 6241–6246.

Shah N, Nute MG, Warnow T, Pop M. 2018. Misunderstood parameter of NCBI BLAST impacts the correctness of bioinformatics workflows. Bioinformatics: bty833.

Singer E, Bushnell B, Coleman-Derr D, Bowman B, Bowers RM, Levy A, Gies EA, Cheng J-F, Copeland A, Klenk H-P, et al. 2016. High-resolution phylogenetic microbial community profiling. The ISME Journal 10: 2020–2032.

Tedersoo L, Anslan S, Bahram M, Põlme S, Riit T, Liiv I, Kõljalg U, Kisand V, Nilsson H, Hildebrand F, et al. 2015. Shotgun metagenomes and multiple primer pair-barcode combinations of amplicons reveal biases in metabarcoding analyses of fungi. MycoKeys 10: 1–43.

Tedersoo L, Ave T-K, Anslan Sten. 2017a. PacBio metabarcoding of Fungi and other eukaryotes: errors, biases and perspectives. New Phytologist 217: 1370–1385.

Tedersoo L, Bahram M, Põlme S, Kõljalg U, Yorou NS, Wijesundera R, Ruiz LV, Vasco-Palacios AM, Thu PQ, Suija A, et al. 2014. Global diversity and geography of soil fungi. Science 346: 1256688.

Tedersoo L, Bahram M, Puusepp R, Nilsson RH, James TY. 2017b. Novel soil-inhabiting clades fill gaps in the fungal tree of life. Microbiome 5: 42.

Wahl HE, Raudabaugh DB, Bach EM, Bone TS, Luttenton MR, Cichewicz RH, Miller AN. 2018. What lies beneath? Fungal diversity at the bottom of Lake Michigan and Lake Superior. Journal of Great Lakes Research 44: 263–270.

Wang Q, Garrity GM, Tiedje JM, Cole JR. 2007. Naive Bayesian classifier for rapid assignment of rRNA sequences into the new bacterial taxonomy. Applied and Environmental Microbiology 73: 5261–5267.

White TJ, Bruns T, Lee S, Taylor J. 1990. Amplification and direct sequencing of fungal ribosomal RNA genes for phylogenetics. In: PCR Protocols. San Diego: Academic Press, 315–322.

Wijayawardene NN, Pawłowska J, Letcher PM, Kirk PM, Humber RA, Schüßler A, Wrzosek M, Muszewska A, Okrasińska A, Istel Ł, et al. 2018. Notes for genera: basal clades of Fungi (including Aphelidiomycota, Basidiobolomycota, Blastocladiomycota, Calcarisporiellomycota, Caulochytriomycota, Chytridiomycota, Entomophthoromycota, Glomeromycota, Kickxellomycota, Monoblepharomycota, Mortierellomycota, Mucoromycota, Neocallimastigomycota, Olpidiomycota, Rozellomycota and Zoopagomycota). Fungal Diversity 92: 43–129.

Wurzbacher C, Nilsson RH, Rautio M, Peura S. 2017. Poorly known microbial taxa dominate the microbiome of permafrost thaw ponds. The ISME Journal 11: 1938–1941.

Yang C, Schaefer DA, Liu W, Popescu VD, Yang C, Wang X, Wu C, Yu DW. 2016. Higher fungal diversity is correlated with lower CO_2_ emissions from dead wood in a natural forest. Scientific Reports 6: 31066.

Zhang J, Kobert K, Flouri T, Stamatakis A. 2014. PEAR: a fast and accurate Illumina Paired-End reAd mergeR. Bioinformatics 30: 614–620.

