## Supplemental Figures for "Combining the 5.8S and ITS2 gene regions to improve classification of fungi"

**Fig. S1** Classification accuracy with ITS2 at different taxonomic levels with different values for the stringency parameter.

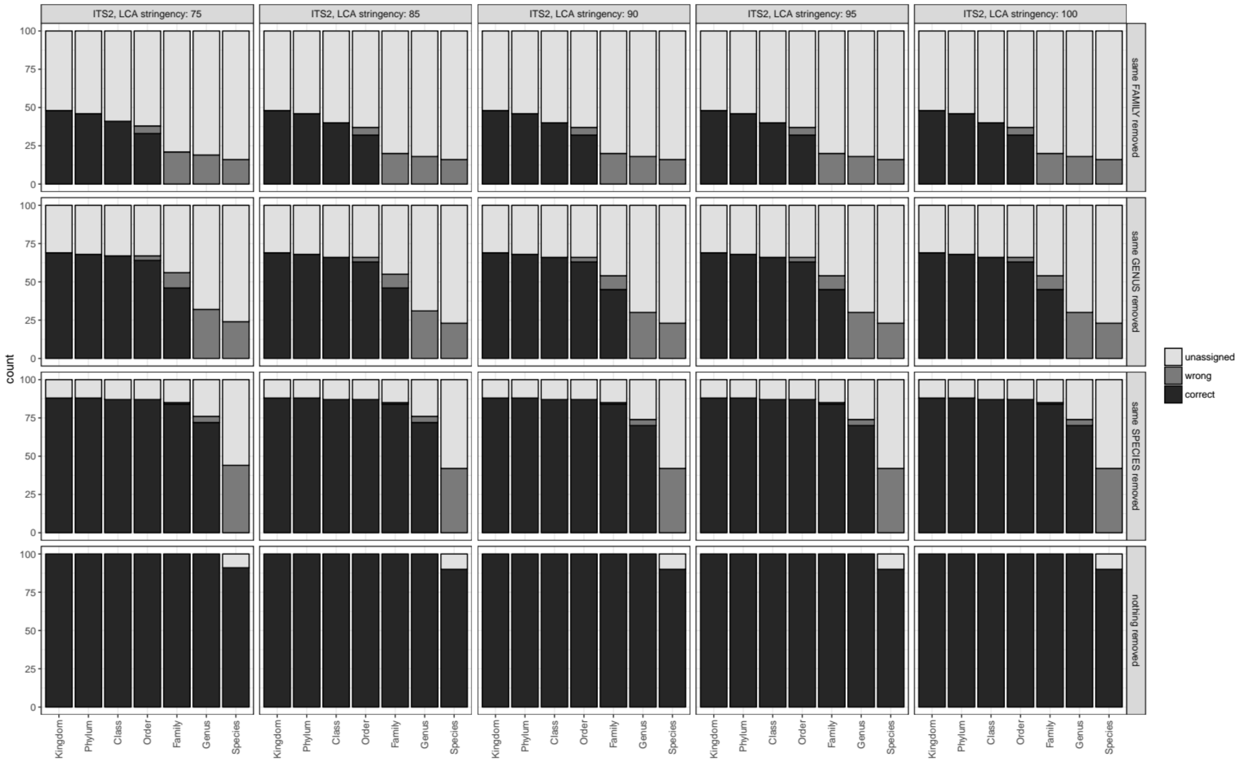

**Fig. S2** Classification accuracy with ITS2 at different taxonomic levels with different values for the minimum identity parameter.

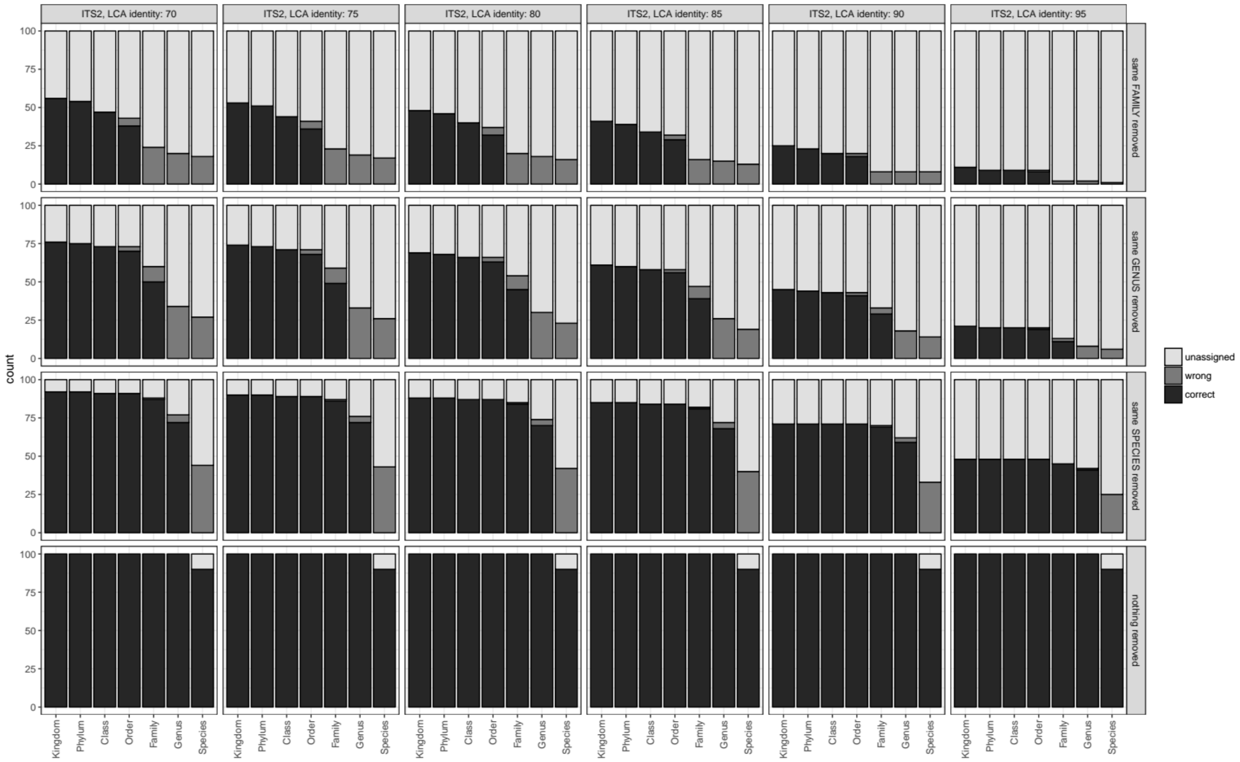

**Fig. S3** Classification accuracy with ITS2 at different taxonomic levels with different values for the minimum coverage parameter.

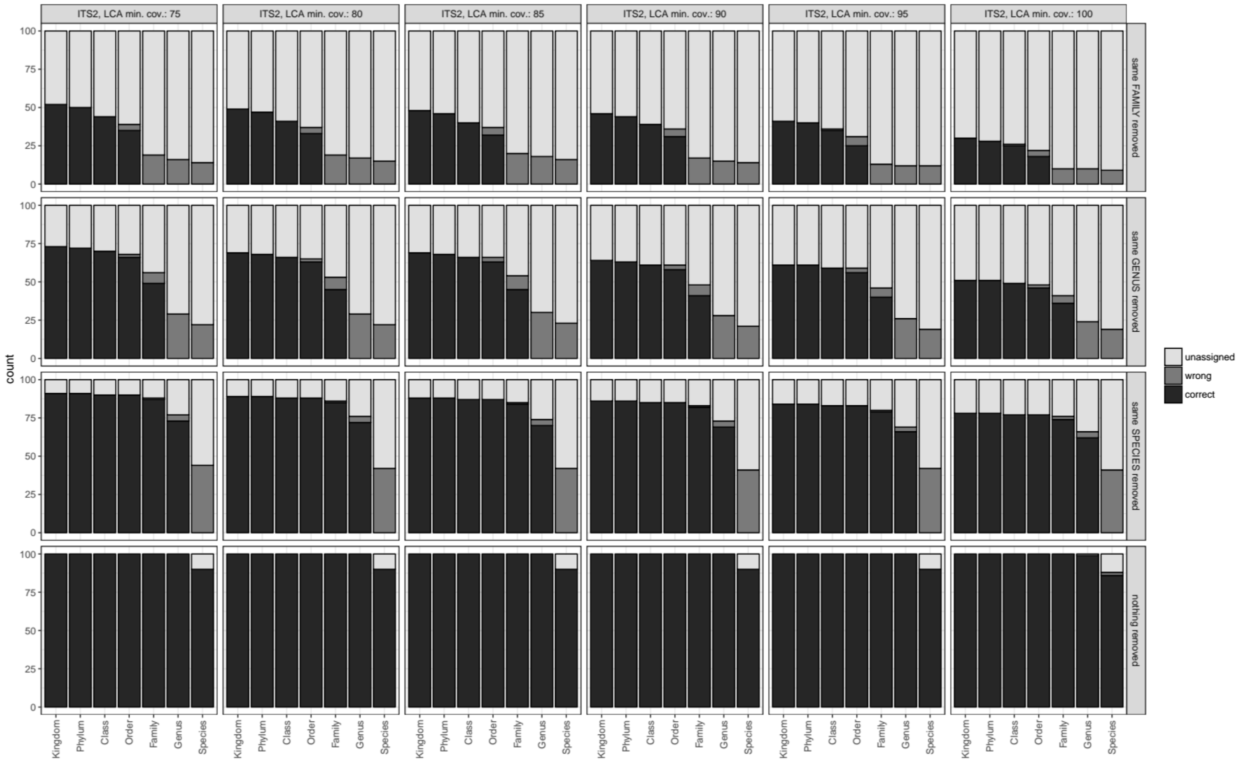

**Fig. S4** Classification accuracy with ITS2 at different taxonomic levels with different values for the top-percent parameter.

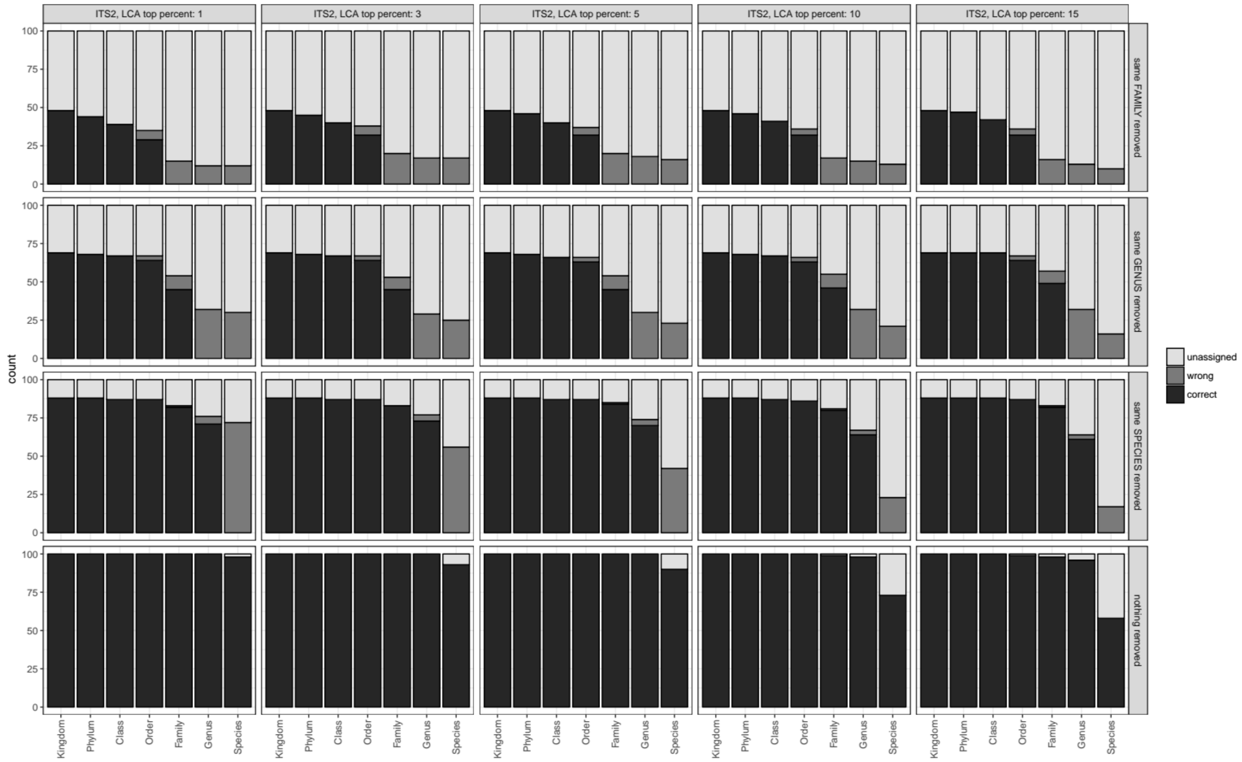

**Fig. S5** Classification accuracy with 5.8S at different taxonomic levels with different values for the identity parameter.

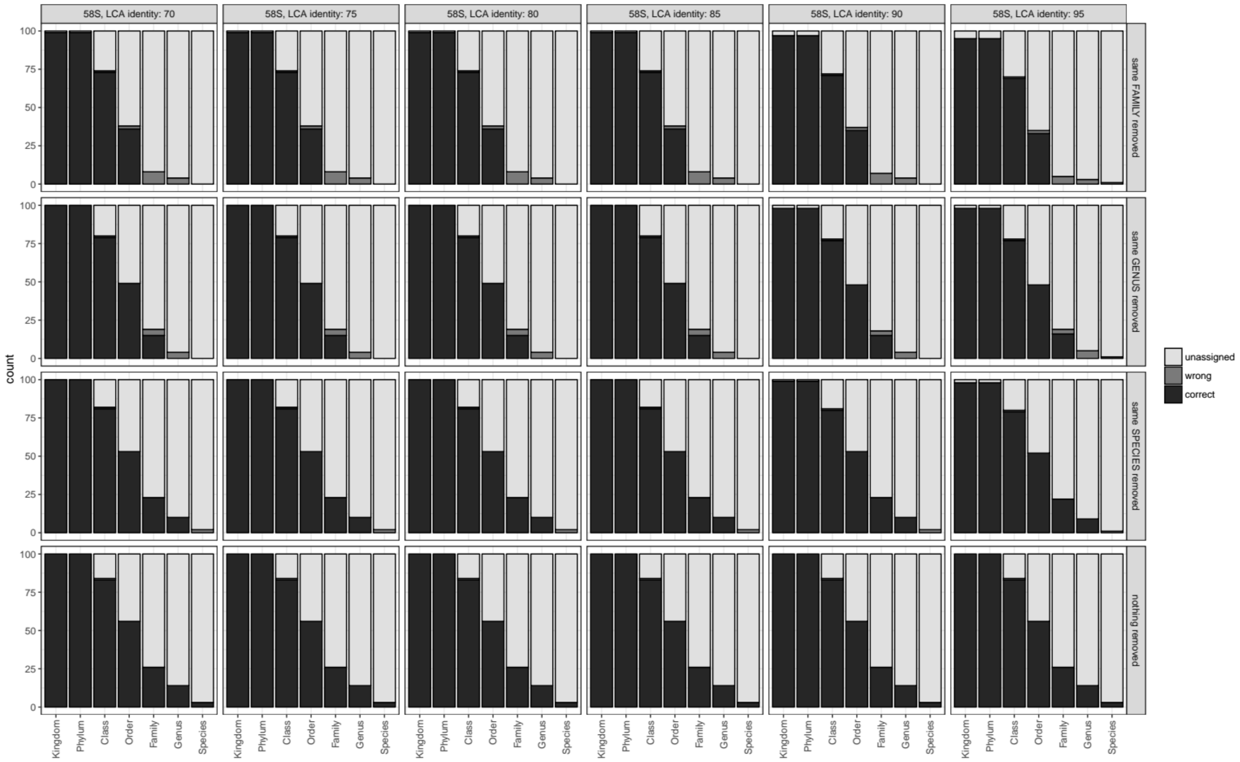

**Fig. S6** Classification accuracy with 5.8S at different taxonomic levels with different values for the minimum coverage parameter.

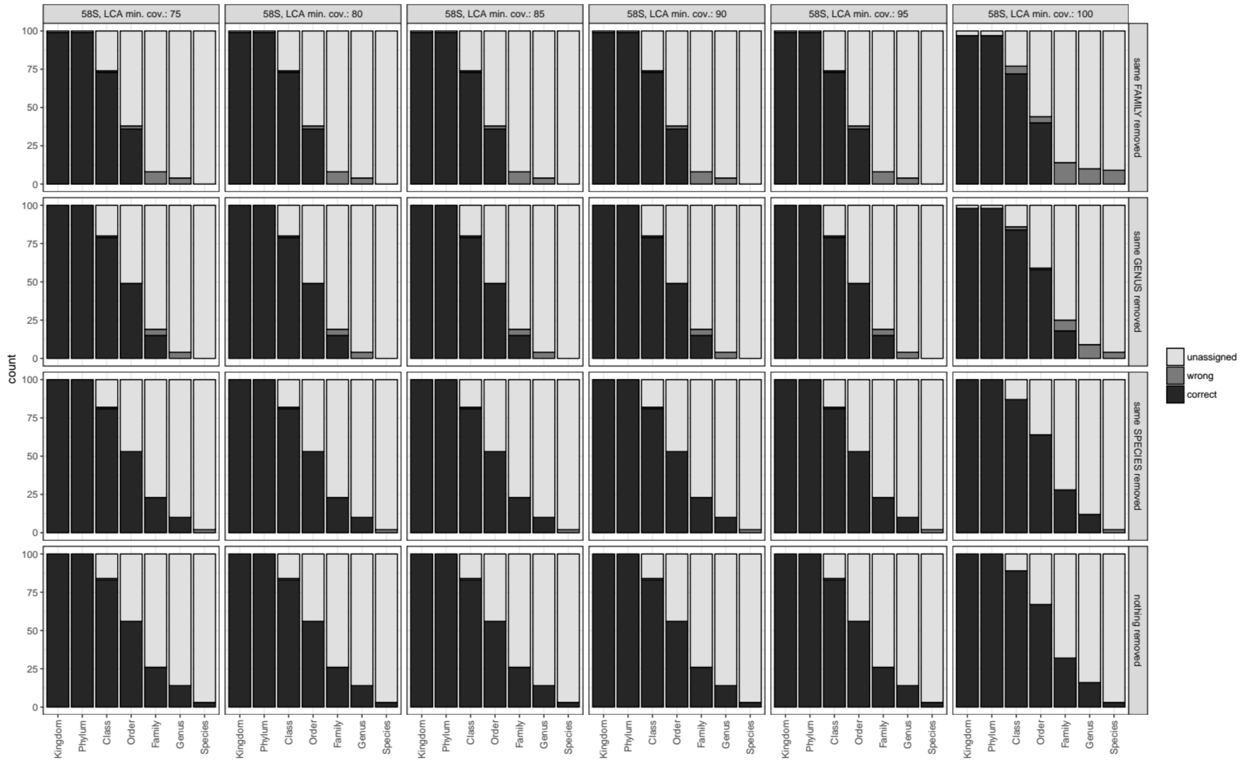

**Fig. S7** Classification accuracy with 5.8S at different taxonomic levels with different values for the top-percent parameter.

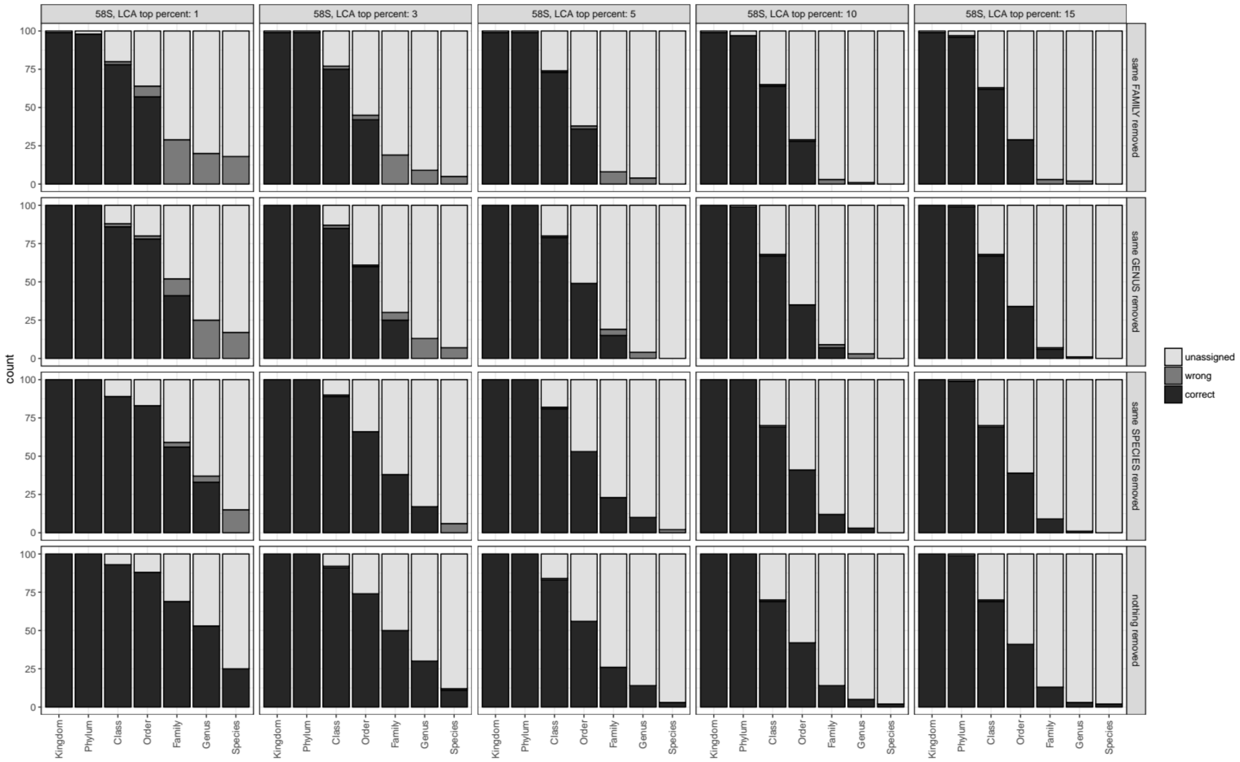

**Fig. S8** Classification accuracy with 5.8S at different taxonomic levels with different values for the stringency parameter.

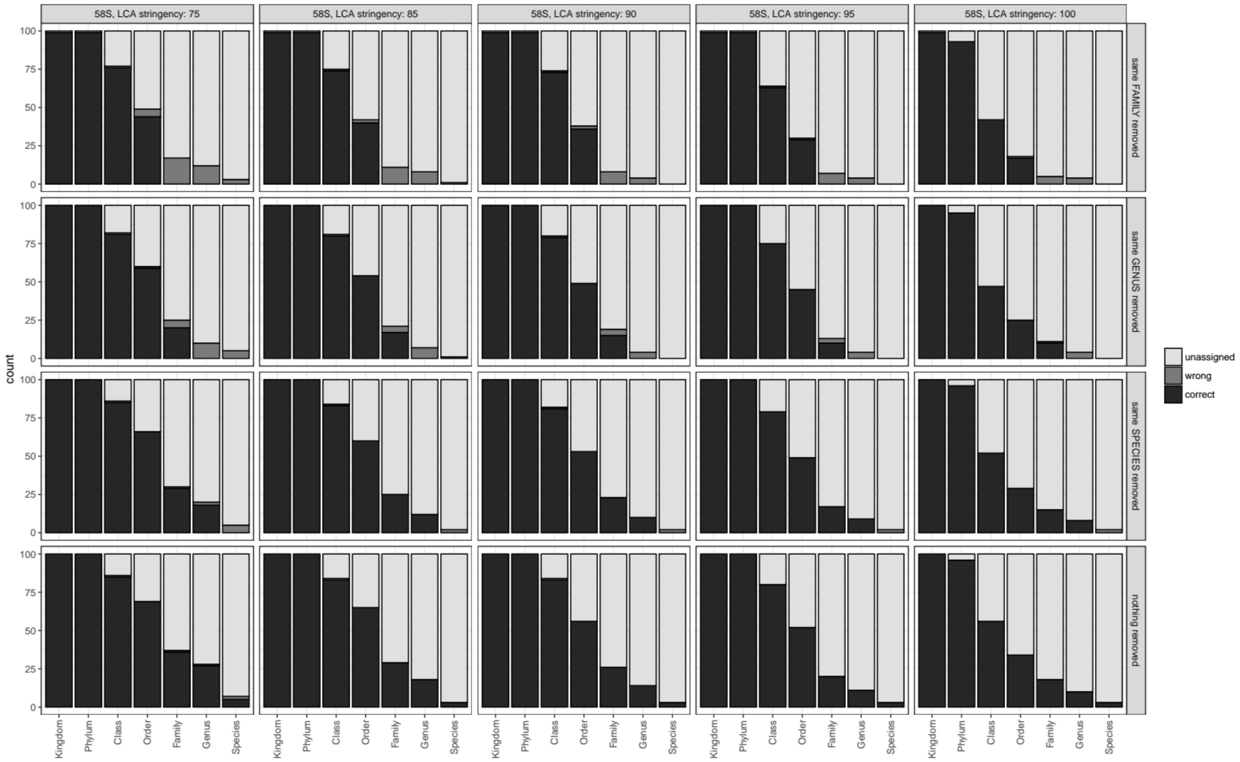

**Fig. S9** Classification accuracy with ITS2 with the LCA approach used in this article, the RDP classifier trained on the UNITE database (RDP_U), and the RDP classifier trained on the Warcup dataset (RDP_W).

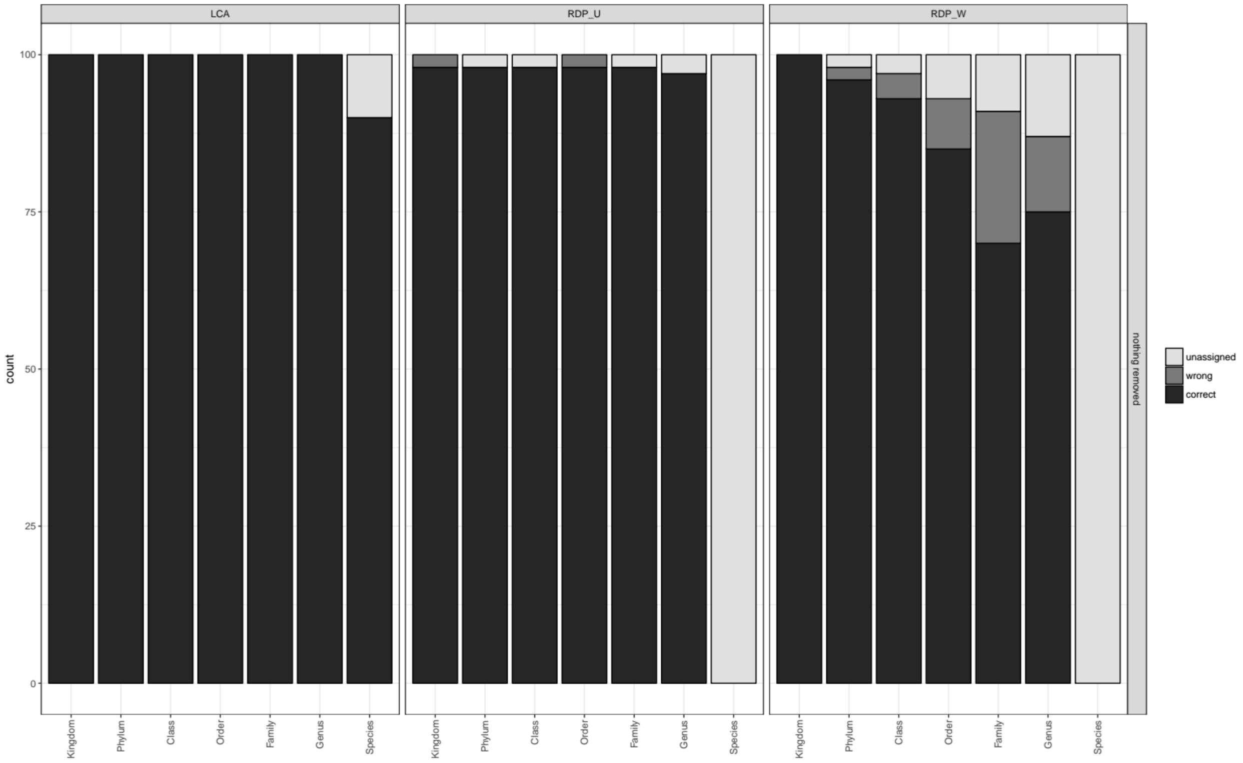

**Fig. S10** Classification accuracy with the combined sequence of 5.8S and ITS2 (left), with 5.8S alone (middle), and with ITS2 alone (right) at different taxonomic levels and with different levels of database completeness.

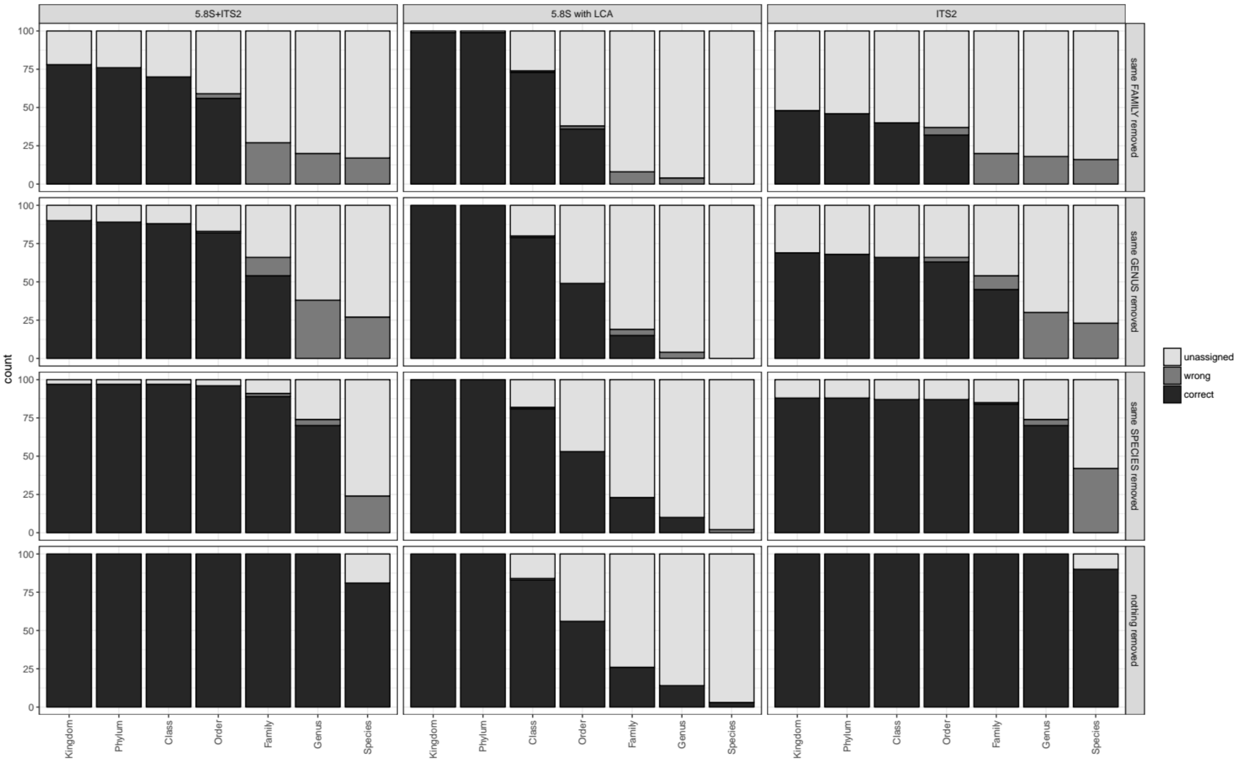
